# When one phenotype is not enough – divergent evolutionary trajectories govern venom variation in a widespread rattlesnake species

**DOI:** 10.1101/413831

**Authors:** Giulia Zancolli, Juan J. Calvete, Michael D. Cardwell, Harry W. Greene, William K. Hayes, Matthew J. Hegarty, Hans-Werner Herrmann, Andrew T. Holycross, Dominic I. Lannutti, John F. Mulley, Libia Sanz, Zachary D. Travis, Joshua R. Whorley, Catharine E. Wüster, Wolfgang Wüster

## Abstract

Understanding the relationship between genome, phenotypic variation, and the ecological pressures that act to maintain that variation, represents a fundamental challenge in evolutionary biology. Functional polymorphisms typically segregate in spatially isolated populations [1, 2] and/or discrete ecological conditions [3-5], whereas dissecting the evolutionary processes involved in adaptive geographic variation across a continuous spatial distribution is much more challenging [6]. Additionally, pleiotropic interactions between genes and phenotype often complicate the identification of specific genotype-phenotype links [7-8], and thus of the selective pressures acting on them. Animal venoms are ideal systems to overcome these constraints: they are complex and variable, yet easily quantifiable molecular phenotypes with a clear function and a direct link to both genome and fitness [9]. Here, we use dense and widespread population-level sampling of the Mohave rattlesnake, *Crotalus scutulatus*, and show that genomic structural variation at multiple loci underlies extreme geographic variation in venom composition, which is maintained despite extensive gene flow. Unexpectedly, selection for diet does not explain venom variation, contrary to the dominant paradigm of venom evolution, and neither does neutral population structure caused by past vicariance. Instead, different toxin genes correlate with distinct environmental factors, suggesting that divergent selective pressures can act on individual loci independently of their genomic proximity or co-expression patterns. Local-scale spatial heterogeneity thus appears to maintain a remarkably ancient complex of molecular phenotypes, which have been retained in populations that diverged more than 1.5-2 MYA, representing an exceptional case of long-term structural polymorphism. These results emphasize how the interplay between genomic architecture and spatial heterogeneity in selective pressures may facilitate the retention of functional polymorphisms of an adaptive phenotype.

## RESULTS AND DISCUSSION

Rattlesnake (*Crotalus*) venoms are among the most complex exocrine secretions in nature, with tens to hundreds of individual components. They display a puzzling phenotypic dichotomy, with two largely mutually exclusive venom strategies: highly lethal type A venoms, characterized by neurotoxic dimeric phospholipase A_2_, e.g. Mojave toxin (MTX), and less toxic type B venoms, which lack MTX, but are rich in snake venom metalloproteinases (SVMPs) with haemorrhagic activity [10]. Both types seem randomly distributed across the phylogeny of rattlesnakes, and sometimes co-occur within populations of a single species. The Mohave rattlesnake, *Crotalus scutulatus*, is well-known for displaying both venom strategies across a continuous distributional range in the North American deserts (Figure 1A) [11-13]. This species thus represents an ideal system to investigate the evolutionary processes and molecular mechanisms shaping intraspecific phenotypic variation and adaptation within a common genetic background.

**Figure 1.**
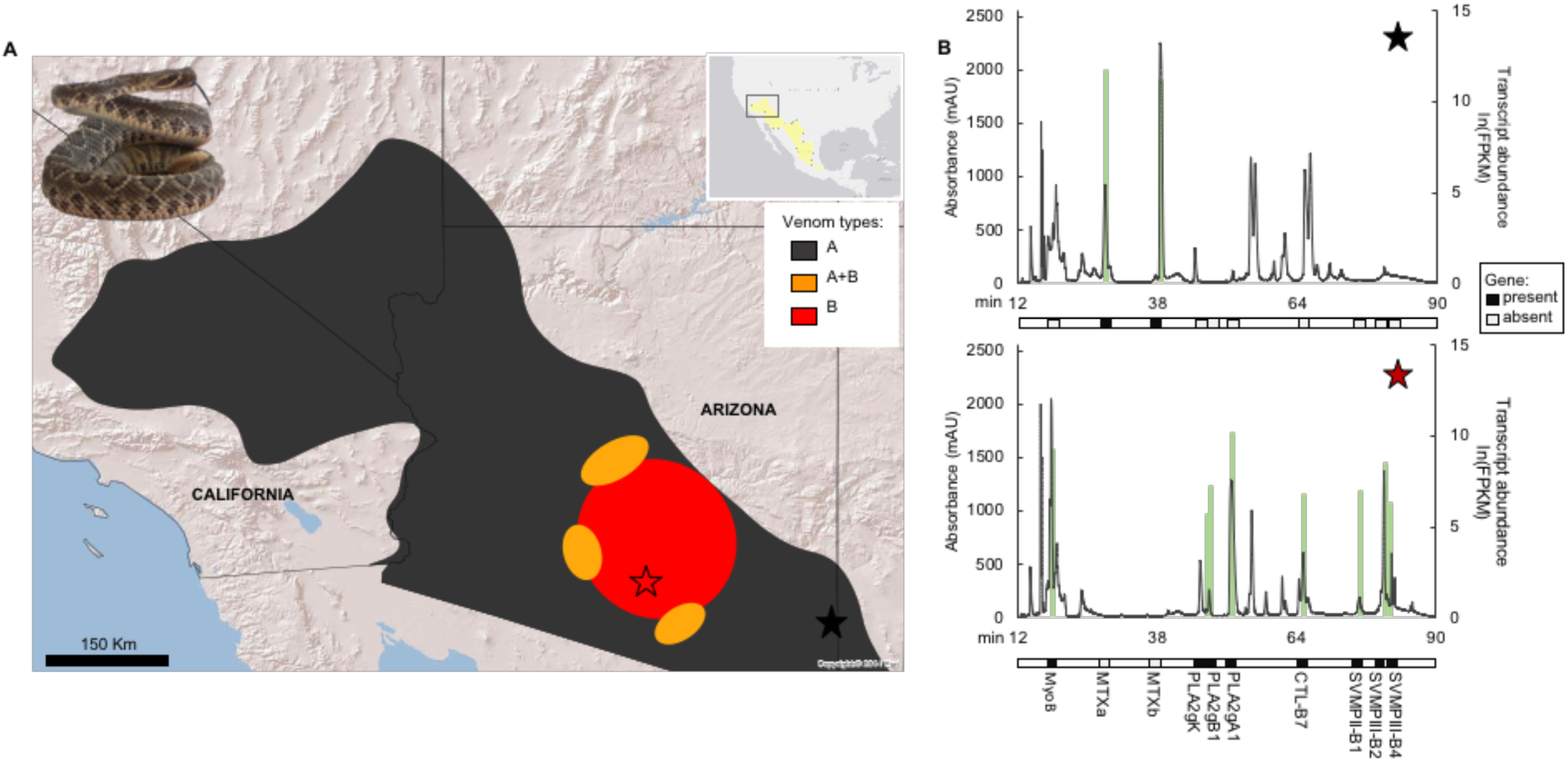
Venom types in Mohave rattlesnakes. (**A**) Distribution of the Sonoran clade of *Crotalus scutulatus* and its major venom types across the study area in the southwestern USA, and sampling location of the two representative individuals. (**B**) Crude venom RP-HPLC chromatograms (black) and transcriptomic expression (green) from type A and B representative individuals with presence/absence of the corresponding genes.

To uncover the mechanisms generating and maintaining polymorphisms across this species, we performed the first densely sampled population-level analysis of the genomic basis of its venom, and used in-depth ecological association analysis (EAA) and climate reconstruction to disentangle the dynamics between genotype, phenotype and environment.

### Venom variation is due to structural genomic variation

To link genotype and phenotype, we generated draft whole genomes, venom-gland transcriptomes and proteomes from one venom A and one type B individual of *C. scutulatus* (ure 1B and S1). As expected, the neurotoxic MTX was highly expressed only in venom A whereas SVMPs were highly expressed only in venom B, but the expression of other toxin genes, including PLA_2_s, SVMPs, serine proteases (SVSPs), C-type lectins (CTLs) and myotoxin (MYO), also differed. To understand whether differences at the genome level, i.e. presence or absence of coding genes, explain differential expression of venom proteins in the phenotypes at the population level, as previously documented for MTX and SVMPs [14-15], we compared genomic and venom proteome profiles of 50 individuals from multiple localities, and PCR-amplified 14 toxin genes belonging to five gene families (Table S1). We discovered that proteomic presence or absence of individual toxins was invariably associated with presence or absence of the coding genes. Based on this strict phenotype-genotype link (Figure S1), we analyzed the spatial distribution of toxin genes in a larger sample to identify gene complexes and linkage patterns (Figure 2). In both main venom types, some genes appear tightly linked, whereas others vary independently. In the core venom B area there were two main genotypes, both characterized by the presence of SVMPs, PLA_2_s (gA1, gB1 and gK) and CTL-B7, but differing in the presence of myotoxin (MyoB). Much greater diversity was observed across the venom A genotypes, which were all characterized by the tightly linked neurotoxic MTXa and MTXb, the absence of SVMPs, PLA_2_gK and gB1, but showing great variation in the occurrence of PLA_2_gA1, MyoB and, to a lesser extent, CTL-B7, each with unique spatial distribution patterns. Linkages between gene complexes were disrupted across the contact zone between venom types, where mixed (A+B) genotypes and multiple different gene combinations occur. Interestingly, the intergrade zones also produced three individuals which possessed neither neurotoxic MTX nor SVMPs genes (venom type O), suggesting that mating between mixed genotypes can disrupt adaptive genomic linkages and can even lead to the complete disappearance of the most important venom components. This leads to the question how these different genomic variants emerged in the population.

**Figure 2.**
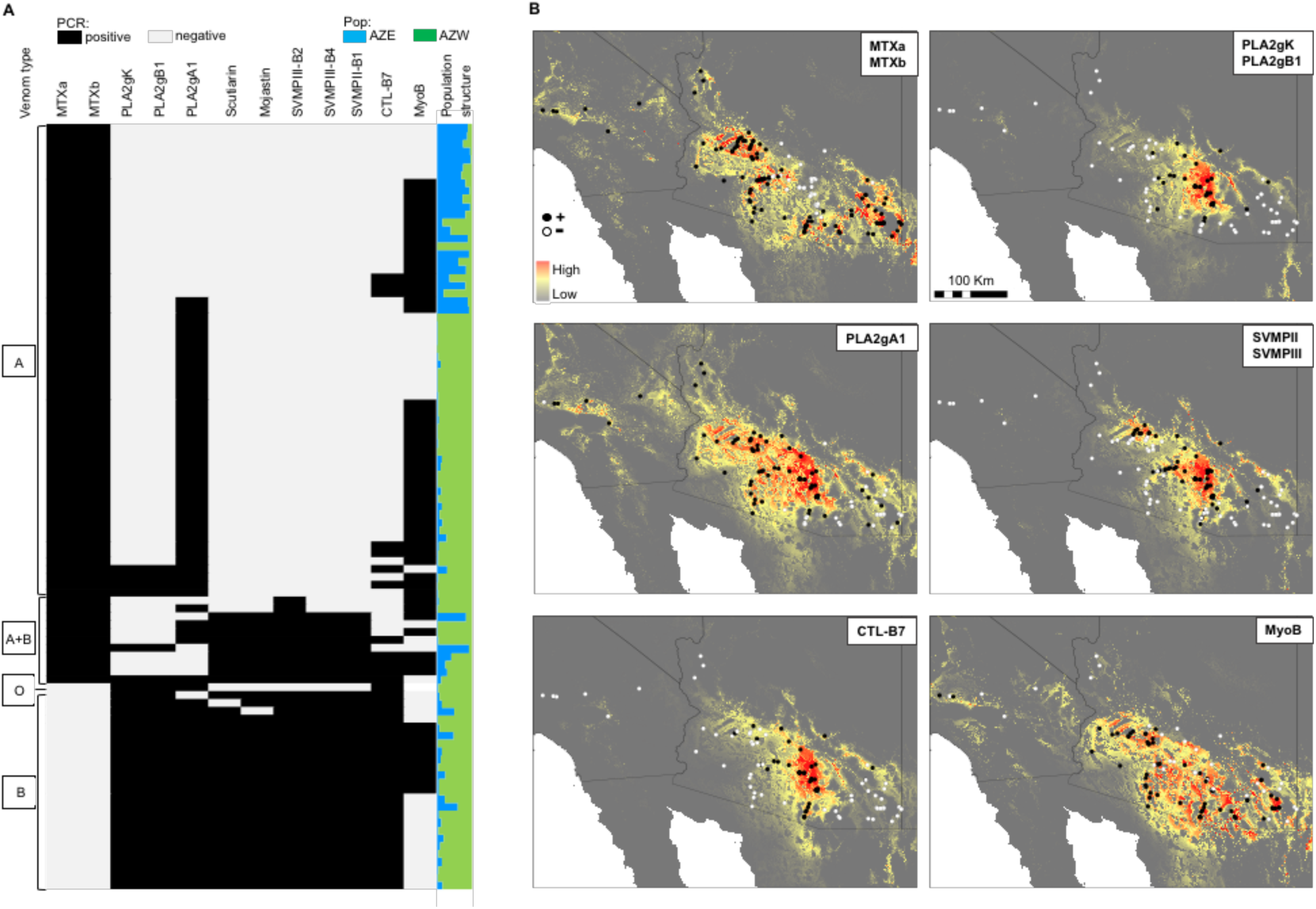
Toxin gene distribution. (**A)** Presence-absence matrix of toxin genes and admixture plot (TESS) with *K*=2. (**B**) Present-day niche models and geographic distribution of toxin genes. Tightly linked genes are shown together.

Most documented recurrent genomic rearrangements [16] and gene family expansions [17] results from non-allelic homologous recombination (NAHR). In rattlesnakes, the diversity at the PLA_2_ gene cluster between three species was previously shown to originate by gene losses from a larger ancestral complex by means of transposable elements (TE)-mediated NAHR [18]. Here, we identified multiple repetitive sequences not only in the PLA_2_ gene cluster, but also in other large, highly variable toxin families, including SVMPs, CTLs, and SVSPs. In these scaffolds, two retrotransposons (ERV and CR1) and one DNA transposon (hAT) were abundant, as in the PLA_2_ cluster where these TEs form conserved blocks in intergenic regions corresponding to the breaking points [18]. The presence of TEs in the genomic scaffolds of multiple polymorphic gene families would support TE-mediated NAHR as a common mechanism causing structural variation, with conserved TE sequence blocks providing the substrate for pairing during recombination resulting in gene content alteration in the progeny cells.

### Venom variation is not associated with population genetic structure

While these results explain the genomic mechanisms underlying the origin of the remarkable venom polymorphism of *C. scutulatus*, the questions of how it persists in the species, and what determines the distribution of venom phenotypes remain. Present-day genetic structure constitutes one plausible hypothesis: populations with high gene flow should have more similar venoms than less connected populations [19]. Alternatively, range fragmentation due to Pleistocene climatic cycles could underlie the spatial segregation of structural variants and venom divergence [20]: fixation of structural variants and gene complexes could have accumulated in vicariant populations, giving rise to intergrade zones after secondary contact following range expansion. Climatic niche modelling of the Sonoran desert clade of this species [21] and reconstruction of its likely distribution through climatic cycles predict that, during the last glacial maximum (LGM), the range contracted into two isolated refugia: an eastern, Madrean population, and a western, Sonoran population (Figure S2). Population genetic analysis of 16 unlinked neutral microsatellite loci identified two genetic clusters with extensive admixture and evidence of post-LGM range expansion (Table S2), which reflect the predicted Pleistocene vicariance. However, the spatial distribution of venom types does not correspond to these genetic clusters. Thus, our results reject the hypothesis of past isolation as a cause for the venom dichotomy in this species.

We further assessed the relationship between venom composition and genetic structure by grouping the samples geographically into localities (Figure S3) and calculating venom distance matrices and toxin gene frequencies. Overall genetic differentiation was weak, including between venom A and B localities (Fst = 0.003-0.05) with high levels of gene flow (Nm = 8-75). AMOVA analysis grouping either by venom types or localities confirmed an absence of finer substructure, with most of the variance arising from within individuals (Table S3). Partial Mantel tests showed weak association between venom variation and neutral genetic distance; similarly, toxin gene frequencies were not correlated with gene flow (Table 1). The complete absence of association between phenotypic variation and neutral genetic differentiation suggests that strong selective forces independent of gene flow are driving the distribution of venom types, not past vicariance.

**Table 1.**
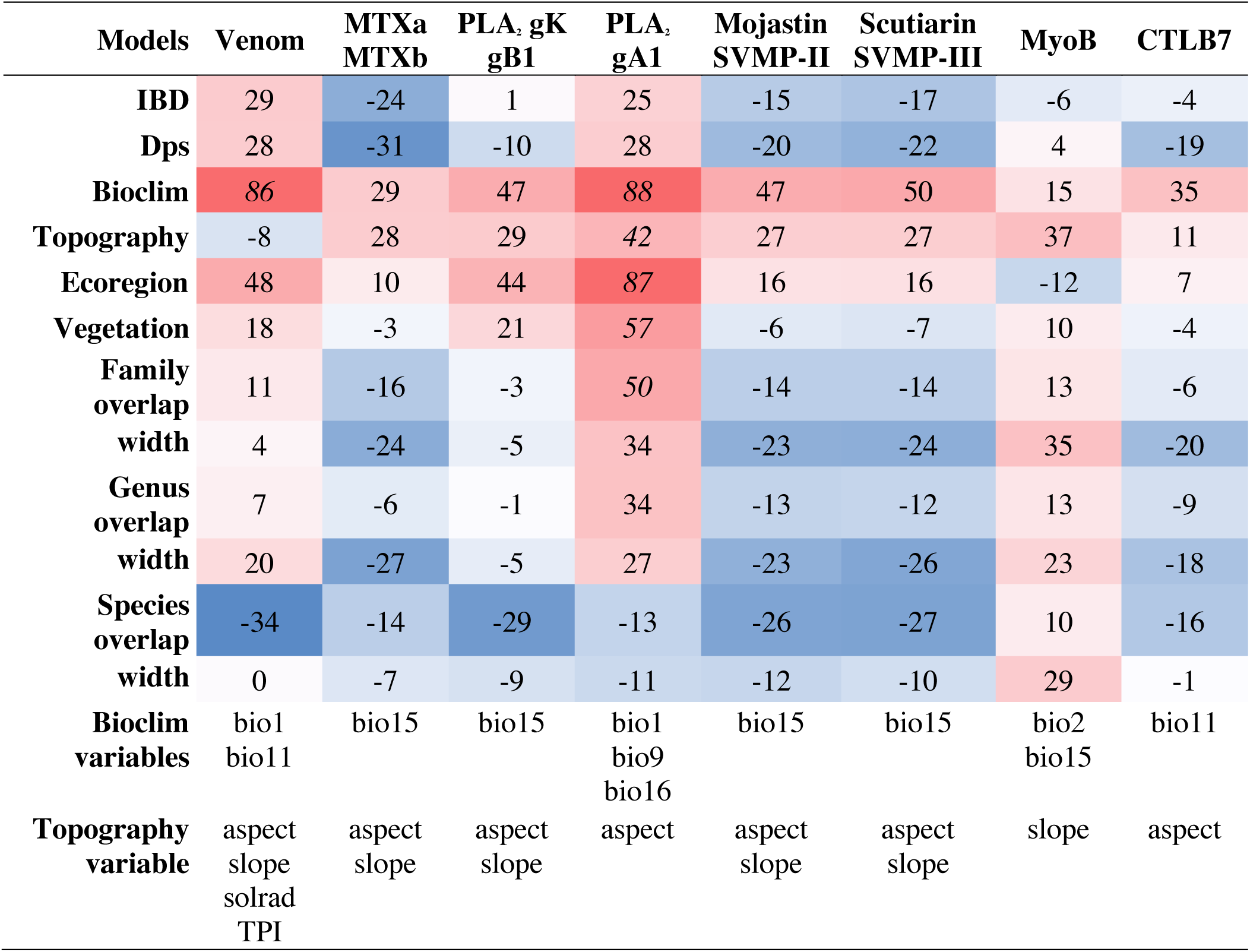
Environmental association analysis between localities. Correlation matrix of partial Mantel tests between venom phenotype or individual toxin gene frequencies (dependent variables) against environmental variables, with geographic Euclidean distance matrix included as covariate. Isolation by distance (IBD) is the null model. Numbers shown in the heatmap are Spearman *R* partial correlation coefficients multiplied by 100. Proportion of shared alleles (Dps) was used as index for neutral genetic differentiation. Variables selected with the BIOENV procedure to generate climatic and topography distance matrix are reported. Prey spectrum was tested at three taxonomic levels using two metrics: overlap (Bray-Curtis), and niche width (Shannon diversity index). Values with p<0.05 are in *italics*. Gene frequencies were not significantly correlated with either diet composition or niche width, except PLA2gA1 which showed a significant negative correlation with diet width (family level: r2= 0.49, p= 0.02).

### Venom composition is not associated with diet

Because the primary function of venom in snakes is the acquisition of prey (*9*], adaptation to specific diet, or local prey communities is generally invoked as the foremost driver of venom evolution [13, 15, 22-24], and has become the dominant paradigm in the study of snake venom evolution. Since even subtle intraspecific variation in venom composition can reflect selection for local prey [24, 25], we hypothesized that the stark contrast in toxicity and mode of action (neurotoxic vs. haemorrhagic) between A and B venoms in *C. scutulatus* would have a significant impact on the snakes’ foraging biology. We therefore tested whether the divergent phenotypes are associated with differences in local diet. Unexpectedly, there was no significant correlation between overall venom variation and similarity in diet niche at any of the taxonomic levels investigated (Table 1).

Similarly, we found no significant pairwise relationships between frequencies of toxin genes and individual prey species; in particular, neither MTX nor SVMPs, the two main players in the venom dichotomy, were linked to any specific prey. We also tested the hypothesis of an association between toxin diversity and diet niche width, with complex venoms allowing predation upon a more diverse array of prey than more streamlined venoms [26]. Interestingly, we found the opposite trend, with venoms A less diverse than B but correlated with broader prey spectra (Figure S3). None of the frequencies of the individual toxin genes were significantly correlated with either diet composition or niche width, except PLA_2_gA1 which showed a significant negative correlation with diet width (family level: *r*^*2*^= 0.49, p= 0.02). Our results thus refute the widespread assumption of diet composition as the main determinant of the venom dichotomy in rattlesnakes, and its universality as a selective driver of snake venom evolution in general [9]. This finding is especially surprising in the light of the apparent role of diet as a driver of much more subtle variation in venom composition in other rattlesnake species [24-25]

### Spatial environmental heterogeneity predicts venom variation

Spatial heterogeneity in environmental variables is a key driver of genotypic and phenotypic polymorphism [27]. In the absence of a strong venom-diet association, we performed EAA to understand whether differences in other biotic and/or abiotic factors contribute to geographic variation of venom composition and toxin genes [28]. Overall venom variation was strongly associated with temperature (Table 1), and the longitudinal climatic gradient characterizing the Sonoran desert (Figure S4) was reflected in the differentiation across venom A profiles (Figure 3A). Yet, the divergence between A and B venoms was not strongly correlated with any specific environmental variable (Table S4). However, across a large, continuous distribution without discrete physical barriers, large-scale analyses may fail to detect the effect of local ecotones and short environmental clines of potential selective importance. We thus analyzed local scale climatic trends along two A-B transects and discovered the presence of sharp clines associated with venom composition for several ecological variables, especially those related to precipitation (Figure 3B-G). While it is unlikely that climate, per se, directly influences venom composition, climatic stability and seasonality may profoundly affect other factors, for instance, prey community dynamics. These, in turn, could influence snake foraging strategies, and potentially also the exposure of snakes to predation, an understudied source of selection on venom [29]. In a widely and continuously distributed species such as *C. scutulatus*, which occupies a variety of environmental conditions, spatial heterogeneity can thus impose selective pressures for local fitness optima, resulting in the maintenance of disparate, locally adaptive gene complexes.

**Figure 3.**
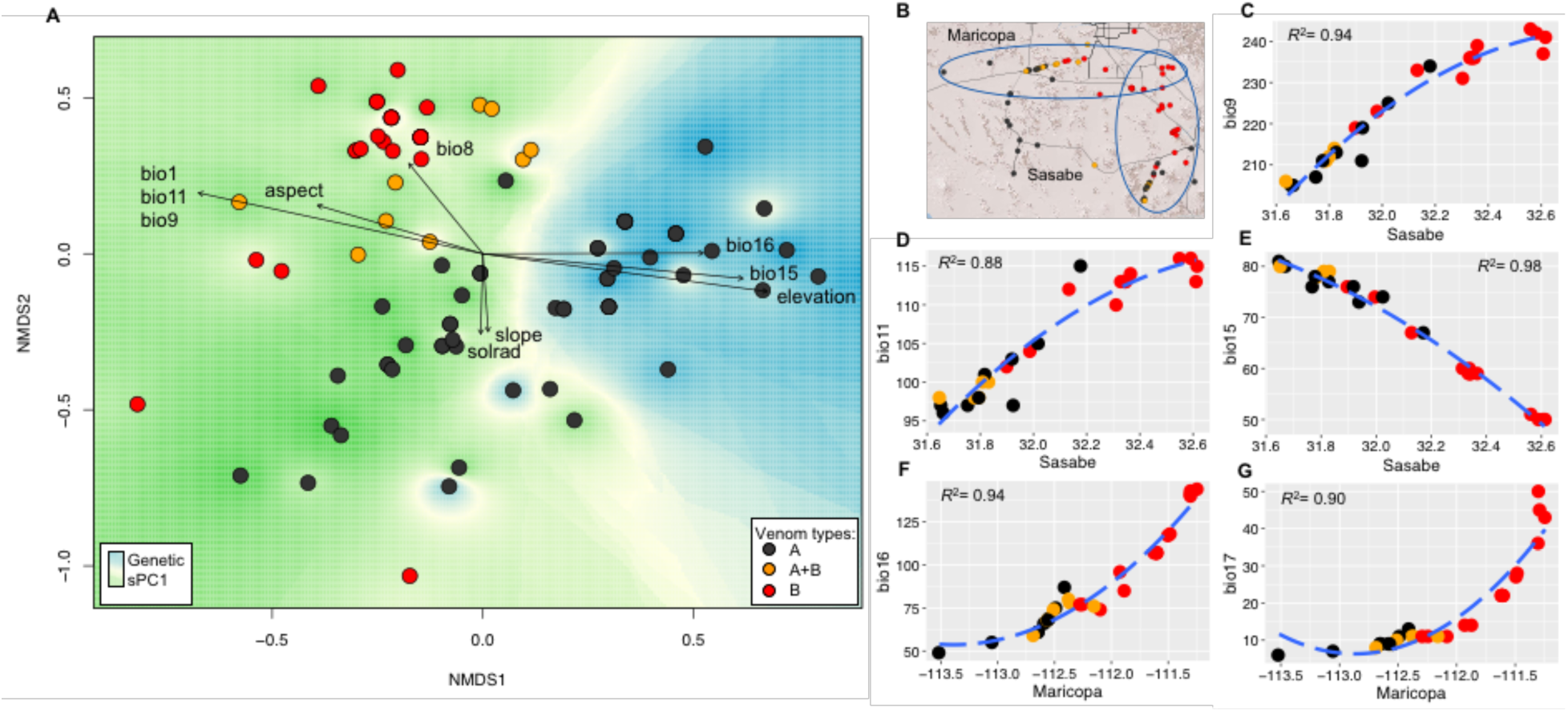
Association between venom phenotypic variation, neutral genetic differentiation and environment. (**A**) Non-metric multidimentional scaling (NMDS) analysis of venom profiles shows great overall variation. The A-B dichotomy is discordant with neutral genetic differentiation (spatial PCA of neutral genotype data). Variation along NMDS1 is strongly correlated with the marked east-west environmental cline across Arizona (Figure S4), whereas environmental associations along NMDS2, corresponding to the A-B transition, are weaker because global-scale variation hinders the detection of local-scale patterns (Table S4). (**B** to **G**) Local-scale analysis along two transects (B) reveals sharp clines in temperature (C-E) and precipitation (F, G) across the venom A-B transition zone.

Logistic regression models revealed that certain toxin genes, even though highly co-expressed in some phenotypes, correlate with different ecological variables, suggesting that multiple selective forces orchestrate individual loci to create complex, dynamic phenotypes (Table S4). Strikingly, genes located few kb apart, such as the PLA_2_s [18], also display independent associations, demonstrating that divergent selective pressures can differentially affect parts of the same genomic region. This is a novel mechanism, considering that genes coding for the same adaptive phenotype are generally brought closer together by means of chromosomal rearrangements such as inversions or supergenes [30].

Another distinctive feature of the rattlesnake venom system is the retention of structural polymorphisms over evolutionary timescales; multiple venom phenotypes are also found in the Chihuahuan clade of *C. scutulatus* [31], suggesting that this structural variation predates the separation of these clades approximately 1.5-2.1 Mya. This multilocus architecture thus represents an exceptional case of long-term maintenance of ancestral polymorphism.

### Genome, environment and the maintenance of geographic variation

The emerging picture of the mechanisms and drivers governing venom variation in *C. scutulatus* is thus one of an ancient retained ancestral polymorphism, with the distribution of toxin genes shaped by natural selection for environmental factors other than diet or neutral gene flow. Disruptive selection against intermediate A+B phenotypes ensures spatial segregation, thereby favoring persistence of gene complexes and divergent phenotypes. The role of relatively subtle environmental changes in driving the dramatic differences in venom composition in this species, coupled with selection against intermediate phenotypes, suggests the existence of “tipping points”, where one phenotype gains a selective advantage over the other.

The unique genomic and genetic architecture of rattlesnake venom also provides important additions to the catalog of mechanisms implicated in driving phenotypic changes in adaptive traits, and establishes a promising system for understanding the ecological and evolutionary implications of genomic structural variation in non-model organisms. Together, our results emphasise the importance of combining large-scale genotype, phenotype and ecological data in natural populations to uncover the wide variety of mechanisms and drivers underlying phenotypic variation, and caution against the rapid acceptance of paradigms as universal, based on a very limited number of case studies.

## MATERIALS AND METHODS

### Model system

Variation of venom composition in rattlesnakes revolves around a dichotomy between two mutually exclusive phenotypes: venom type B, characterized by high expression levels of snake venom metalloproteinases (SVMP) with hemorrhagic and proteolytic activity, and venom type A, rich in a highly lethal presynaptic β-neurotoxic heterodimeric phospholipase A_2_ (PLA_2_) (e.g. Crotoxin, Mojave toxin - MTX) but lacking SVMPs [10]. The distribution of these venom phenotypes across the phylogeny of rattlesnakes is highly irregular, with most clades having representatives secreting both venom types. Moreover, some rattlesnake species possess both phenotypes with individuals secreting either type A or B venoms.

The Mohave rattlesnake (*Crotalus scutulatus*), a widespread species found across the southwestern USA and Mexico, represents an ideal system to study the causes and mechanisms underlying variation in this remarkable molecular phenotype. Three distinct phylogeographic lineages have been identified: eastern Mexico, including the subspecies *C. s. salvini,* the Chihuahuan desert and the Sonoran desert (which includes the Madrean Archipelago and Mojave desert) [21]. Populations of all three clades in much of the US and southern parts of the Mexican range secrete type A venoms, whereas in central Arizona as well as northern and central parts of Mexico, Mohave rattlesnakes secrete type B venoms [11-13, 31]. Intermediate A+B venoms containing both SVMPs and MTX are found at the contact zones between the two venom types. Besides SVMPs and the neurotoxic PLA2, additional toxins belonging to different gene families show geographic variation in their expression, in particular myotoxin, a potent voltage-gated potassium channel inhibitor which causes rapid paralysis and muscle necrosis [12, 13].

### Sample collection

*Crotalus scutulatus* venom and blood or tissue samples were obtained from field-caught specimens from most of the distribution of the Sonoran clade [21] from California to south-western New Mexico, as well as from western Texas. All field-collected specimens were released unharmed at the locality of capture within 72h. Two field-caught adult snakes, one male from venom A (N31.85791, W109.04778) and one female from venom B areas (N31.90326, W111.39112), were temporarily housed at Bangor University prior to venom-gland dissection. These specimens were chosen as representative for in-depth proteomic, transcriptomic and genomic analyses. The two representative snakes were tested for presence of Mojave toxin (MTX) genes to correctly assign them as either venom A or B.

### Venom proteomics

Crude dried venoms of 50 adult snakes were dissolved in 0.05% trifluoroacetic acid (TFA) and 5% acetonitrile, and insoluble material was removed by centrifugation prior to reverse-phase HPLC. Proteins were separated on an Agilent 1100 system using a Teknokroma Europa 300 C18 column (250 × 4 mm, 5 μm particle size) eluting at 1 mL/min with a linear gradient of 0.1% TFA in water and acetonitrile, and chromatographic fractions were manually collected. Molecular masses of RP-HPLC purified proteins of the two representative venom types and a subset of specimen from different geographic areas were determined by electrospray ionization (ESI) mass spectrometry using an Applied Biosystems QTrap™ 2000 mass spectrometer operated in Enhanced Multiple Charge mode in the range m/z 350-1700, and data acquired and processed using Analyst 1.5.1. software. Chromatographic fractions were estimated by SDS-PAGE on 15% polyacrylamide gel and protein bands excised and subjected to automated in-gel digestion using a ProGest Protein Digestion Workstation (Genomics Solutions Ltd.) and sequencing-grade porcine trypsin (Promega). Tryptic digests were dried in a SpeedVac (Savant™, Thermo Scientific Inc.), re-dissolved in 15 μL of 5% acetonitrile containing 0.1% formic acid, and submitted to LC-MS/MS. Tryptic peptides were separated by nano-Acquity UltraPerformance LC^®^ (UPLC^®^, Waters Corporation) using BEH130 C18 (100 μm × 100 mm, 1.7 μm) column in-line with a Waters SYNAPT G2 High Definition Mass Spectrometry System (Waters Corporation). Doubly- and triply-charged ions were selected for collision-induced dissociation (CID) MS/MS. Fragmentation spectra were interpreted manually or using the on-line form of the MASCOT program (http://www.matrixscience.com) against the NCBI non-redundant database. Additionally, all peptide sequences were blasted against the venom-gland transcriptome assemblies using tblastn adjusted for short sequences in order to directly link venom proteins to their corresponding transcripts.

Presence or absence of MTX and SVMP peaks in the RP-HLPC chromatograms were used to assign individuals to either venom type A, B or A+B for further analyses.

### Venom fingerprinting

To compare venom phenotypic variation and diversity in a consistent and reliable way, we used our previously described method [32] and characterized 98 *C. scutulatus* venom profiles by fingerprinting. The binary matrix obtained by scoring protein peaks as either present or absent was used to calculate Shannon diversity index and pairwise Bray-Curtis dissimilarity matrices for subsequent analyses.

### Venom-gland transcriptomics

Venom glands of the two representative *C. scutulatus* venom A and B were dissected four days after venom extraction when transcription levels reach the maxima [33]. Left and right venom glands were removed and immediately frozen in liquid nitrogen. Total RNA was isolated using the Trizol Plus RNA Purification kit (Life Technologies) with an additional DNase I treatment. cDNA libraries were constructed using the TruSeq RNA Sample Preparation kit v3 (Illumina) with a selected fragment size of 300 bp and sequenced using 125 bp paired-end reads on an Illumina HiSeq2500. Pre-processing of raw fastq files was performed with the Trimmomatic tool [34] and quality was assessed using FastQC tool [35]. The resulting high-quality reads from the two venom-gland samples (N50 = 1293 type A, 1286 type B) were assembled separately using Trinity 2.0.4 [36]. Reads were aligned back to the *de novo* assemblies using Bowtie2 [37] and abundances were estimated with the RSEM tool [38] using the Trinity package. Prior to annotation, the assembled transcripts with FPKM (Fragments Per Kilobase of exon per Million mapped reads) value < 1 were removed and identical sequences were filtered out using CD-HIT [39]. In order to find all possible toxin transcripts, the assemblies were subjected to a blastx analysis against multiple databases including the NCBI nonredundant (nr) protein sequences [40], UniProtKB [41] and a custom database containing only toxin protein sequences. Functional annotation of the assemblies was further performed following the Trinotate pipeline [42]. Finally, to obtain higher quality transcript abundance estimates, all transcripts without annotation to any of the databases were filtered out and transcript abundances recalculated. Homologous toxin transcripts between the two venom types were identified by reciprocal blast analysis and considered homologous if the coding sequences were 99% identical and with minimum 70% sequence coverage.

### Draft whole-genome sequencing

We generated draft whole genomes for the two representative individuals. Genomic DNA was extracted from blood samples using the Qiagen DNA blood and tissue kit. For each individual, two genomic libraries were prepared using the Truseq DNA Sample Preparation kit (Illumina) with insert sizes of 300 bp and 600 bp, and sequenced on one lane of an Illumina HiSeq2500 using 125 bp paired-end reads. Pre-processing of raw reads was performed as for the RNA-Seq data. Sequencing coverage was approximately 8x. High quality paired-end reads were assembled *de novo* using the CLC Genomics Workbench platform v6.5. Pre-assembled contigs were orientated and combined into scaffolds using SSPACE Standard 3.0 [43] which improved contig length in both assemblies (N50 = 4430 type A, 5182 type B).

Genomic scaffolds containing putative toxin genes were identified by mapping all toxin transcripts to both genome assemblies using the GMAP software [44].

Transposable elements (TE) in the genomic scaffolds harboring toxin genes were identified with the RepeatMasker web server [45] using the cross_match search engine run in slow mode (higher sensitivity).

### Toxin gene selection and PCR primer design

The most highly expressed and variable toxins identified in the venom proteomes and transcriptomes were selected as candidates for further investigation. The most highly expressed and variable toxins generally belong to medium to large gene families, and isoforms often differ by only few nucleotides, making paralog-specific primer design challenging. In some multigene families such as SVMP and C-type lectin (CTL), several transcript sequences are identical with the exception of one or a few regions coding for surface-exposed amino acid residues in the mature protein, which interact with the target receptors [46]. For example, four SVMP-II transcripts identified in the venom B transcriptome had identical sequences in the disintegrin domain but were divergent in the metalloproteinase domain; similarly, four transcripts of the SVMP-III family had identical metalloproteinase and disintegrin domains, but several nucleotide differences in the cysteine domain. To design gene-specific primer pairs, for each protein family separately, we first identified exon-intron boundaries by aligning with MUSCLE [47] all transcripts against the longest identified genomic scaffolds (see above) and checked the alignments manually. Additionally, FGENESH [48] was used for gene structure prediction using *Anolis carolinensis* or chicken (“Aves generic”) as organism-specific gene-finding parameters. To ensure primer specificity and reliability of PCR amplification, we designed primer pairs following several criteria:

-multiple primer sets were designed for each selected toxin transcript

-primer annealing sites were designed to correspond to uniquely specific gene sequence regions

-primer pairs were designed to anneal on two adjacent exons and spanning one intron (Exon-Primed Intron-Crossing – EPIC PCR)

-when possible, locus-specific primer sets were designed (see below).

PCR primers were designed using the Primer-BLAST tool [49], and amplification specificity was checked against the two *C. scutulatus* transcriptomes and the NCBI nucleotide database.

In total, 23 PCR primer pairs were designed to amplify 14 toxin isoforms belonging to five multigene families. These include: four PLA2 genes, two Myotoxins (MYO), two CTL, six SVMP-II, and two SVMP-III. For the SVMPs primers were designed both in the metalloproteinase and the disintegrin domains. For SVMP-II it was not possible to design gene-specific primers because of the high similarity between isoforms; instead, primer sets were predicted to amplify two isoforms each; similarly, SVMP-III were predicted to amplify two isoforms each, but those differed only 2nt differences, so we considered them as alleles of the same gene. PCR amplification was performed on a total volume of 15 μl using 2X ReddyMix PCR Master Mix (Thermo Scientific), 0.2 mM of primers and 20 ng of gDNA. Cycling conditions were as follow: an initial denaturation at 94°C for 2 min, followed by 35 cycles of 94°C for 30s, 55°-61°C for 30s, 72°C for 1-1:30 min (depending on amplicon size), and a final extension at 72°C for 5 min. PCR products were verified by Sanger sequencing (Macrogen).

All primer sets were first tested on the representative individuals; finally, 12 toxin genes were selected for further investigation (Table S1). In addition, we used previously designed primers for the acidic (MTXa) and basic (MTXb) subunit genes of Mojave toxin [14]. In total, 98-163 individuals were screened for toxin gene presence, PCR products were checked on 1.5% agarose gel, and a subset were sequenced to verify consistency of primer specificity across the geographic distribution. Toxin gene sequences were blasted against the NCBI nucleotide (nt) and whole-genome shotgun contigs (wgs) databases.

Given the strong correlation between the presence/absence of each toxin in the venom proteome and the corresponding coding gene in the genome, we used genotype information to assign additional individuals without proteomic information (e.g., samples from road killed specimens) to venom type A, B and A+B.

### Population genetic analysis

Initially, 366 *C. scutulatus* individuals were genotyped at 16 polymorphic microsatellite loci (Table S2). Briefly, 3-4 loci were simultaneously amplified using the Qiagen multiplex PCR Master Mix and PCR products were run with the GeneScan™ 600 LIZ size standard (Applied Biosystems) on an ABI 3130xl Genetic Analyzer. Run quality, peak calling and allele size scoring was performed with the GeneMapper 4.0 software (Applied Biosystems). The Mohave rattlesnake is known to hybridize in the wild with the Prairie rattlesnake, *C. viridis*, where the ranges of the two species come into contact in south-western New Mexico [14]. To avoid bias or errors in estimates of population genetic structure and toxin gene distribution, all potential hybrids were removed from the analysis. We genotyped 35 field-caught specimens of *C. viridis* from New Mexico and used the program STRUCTURE 2.3.4 [50, 51] for hybrid detection. Ten independent runs with a burn-in period of 100 000 replications and 1 000 000 MCMC iterations were performed with *K* = 2. The individuals assigned to the *C. scutulatus* genetic cluster with posterior probability < 0.95 were considered as potential hybrids and removed from the dataset. To further reduce the possibility of including backcrossed or undetected hybrids, we applied a stricter rule by excluding all specimens collected in close proximity to the hybrid zone (see [14] for detailed map). After data refinement 313 individuals of *C. scutulatus* individuals were genotyped at 16 polymorphic microsatellite loci to estimate neutral genetic variation and population genetic structure.

Neutral genetic structure was determined using the spatial Bayesian clustering algorithm implemented in TESS 2.3.1 [52]. This program detects genetic barriers or discontinuities in continuous populations based on a hierarchical Markov Random Field (MRF) model whose neighborhood system is obtained from a Voronoi tessellation. To select the optimal number of clusters, we first ran TESS without admixture for *K*_max_ ranging from 1 to 6 for 50 000 sweeps and a burn-in of 10 000. The optimal value of *K* _max_ was determined by plotting the Kmax values against the deviance information criterion (DIC) and considered the values for which the DIC first reached a plateau. Then, we performed 10 independent runs using the CAR model with admixture and including Euclidean geographic distances. Admixture coefficients were estimated using the run with the lowest value of DIC, and individuals were assigned to the genetic cluster with coefficient values > 0.5.

Additionally, we explored spatial patterns of neutral genetic variability with spatial Principal Component Analysis (sPCA) in the R package adegenet 2.1 [53]. This method is an adaptation of PCA that optimizes the variance of the principal components and their spatial autocorrelation. Spatial genetic patterns were visualized by interpolating the lagged principal scores from sPCA analysis using the Inverse Distance Weighted (IDW) technique in ArcMap (ESRI^®^ ArcGIS 10.3) to obtain maps of genetic clines for each principal component.

Presence of null alleles was assessed with FreeNA [54] and Genepop 4.4 [55], and each locus was tested for linkage disequilibrium (LD) and conformity to Hardy-Weinberg Equilibrium. Standard summary statistics were calculated using GenAlex [56]. Three loci (Crti95, CS2340 and Ca290) were highly significant for presence of null alleles and deviated from HWE, hence analyses were first run with the whole set of 16 markers, and then excluding these three loci and results were compared. While genetic structure did not differ when using either the full or reduced datasets and pairwise genetic distances were highly correlated (Mantel *r*^2^ = 0.99, p = 0.001), important differences were observed for the HWE and heterozygosity tests within each locality, hence here we report the results obtained from the reduced dataset.

Partitioning of genetic variation within and across subpopulations as inferred by TESS was examined using analysis of molecular genetic variance (AMOVA) in GenAlex. To test whether spatial genetic patterns and population structure are the results of recent genetic bottlenecks, heterozygosity excess and deficit were tested using the software BOTTLENECK v1.2.02 [57] and Genepop.

Presence of isolation by distance (IBD) was tested by analysing the genetic distances between pairs of individuals as a function of geographic distances in GenAlex. Additionally, a pairwise genetic distance matrix was estimated based on the proportion of shared alleles (*Dps*) [58] between localities and used in a Mantel test against Euclidean geographic distances.

### Statistical analysis workflow

Initial analysis was run for the whole dataset including Texas samples. However, recent phylogeographic analysis has confirmed that *C. scutulatus* in the Sonora and in the Chihuahuan desert are two spatially and genetically distinct populations that have been separated for a long period of time [21], and our microsatellite analysis also support this finding. Consequently, as we were interested into dissecting population-level evolutionary processes, we excluded Chihuahuan population from further analysis.

All statistical analyses were performed in R version 3.4.2 [59] using two approaches. First, we grouped individuals into discrete spatially-defined sampling localities. The criteria used to delineate the localities included gaps in sampling distribution and valley/mountain ridge systems. For this subpopulation-level analysis, individuals falling between localities, and thus difficult to assign to a specific group, were excluded. Although this approach has the drawback of removing samples collected between localities, it can take advantage of population-based association approaches, for instance it allows to test for relationships between phenotype and diet composition. In this population-based approach, we ran Mantel and partial Mantel tests (controlling for geographic distance) from the *vegan* 2.4-4 package [60] using the following response distance matrices:

-venom phenotype: mean pairwise Bray-Curtis dissimilarities between localities calculated using the the on-chip fingerprinting binary matrix;

-venom genotype: pairwise Bray-Curtis dissimilarity matrices based on toxin gene frequencies (one for each gene).

All p-values were corrected for multiple comparisons [61]. In one locality (“Gila”) we were not able to collect venoms, however we included this in the genotype analysis as we did have tissues, and hence genetic information.

Secondly, we used an individual-based approach which has the benefit of taking into account all available information and allow for better detection of association along gradients. For the venom phenotype, we analyzed patterns of variation using non-metric multidimensional scaling (NMDS) based on a pairwise Bray-Curtis distance matrix and used the individual scores on the first two axes as response variables in regression models. For the venom genotype, presence or absence of each toxin gene were used as response variables in logistic regression models using the glm (generalized linear model) function with binomial(link=“logit”) error distribution.

### Environmental datasets

Current climatic data were obtained from the WorldClim 1.4 database (http://www.worldclim.org) at 30 sec resolution [62]. To avoid collinearity, variables which were highly correlated (Pearson’s coefficient |*r*| ≥ 0.8) were pruned based on a pairwise correlation matrix leaving a total of 13 climatic variables. Past climatic data for the Mid Holocene (Mid Hol) and Last Glacial Maximum (LGM) were obtained from simulations with Global Climate Models (GCMs) estimated by the Community Climate System Models (CCSM), and data from the Last Interglacial (LIG) were generated by Otto-Bliesner *et al*. [63].

A high resolution digital elevation model (DEM) was obtained from ASTER (http://asterweb.jpl.nasa.gov). The DEM raster was used to produce additional topographic variables including slope, solar radiation, aspect and topographic position index (TPI) using the Spatial Analyst toolbox in ArcMap 10.3 (ESRI^®^).

Land cover data describing North American ecological areas (level III “ecoregions”) was obtained from the US EPA (https://www.epa.gov/eco-research/ecoregions-north-america), and vegetation data were obtained from the Gap Analysis Project (https://gapanalysis.usgs.gov/gaplandcover/data/download/).

Spatial patterns of environmental heterogeneity across the study areas were examined by ordination techniques using Principal Component Analysis (PCA), and significant differences between localities were tested with pairwise t-tests.

### Venom variation and past climatic changes

To test whether present-day phenotypic variation in venom composition is the result of past range fragmentation due to climatic changes, we performed niche modelling using the program MAxEnt [64]. Georeferenced occurrence localities of *C. scutulatus* were gathered from the VertNet (http://vertnet.org) and Global Biodiversity Information Facility (www.gbif.org) online databases and verified for possible mislabeled coordinates prior to the analysis. All models were run with default regularization and 10 replicates subsampled, using 20% of the points for test and 80% for training each replicate. We generated ecological niche models for the species as well as for each individual toxin gene, and used present-day climate envelopes for prediction of past scenarios.

### Venom variation and current gene flow

Multiple approaches were used to test whether variation in venom composition reflects current patterns of gene flow and neutral genetic structure. First, we used the AMOVA procedure in GenAlex to estimate number of migrants and compare molecular variance between (i) the three major venom types (i.e. A, B, A+B), and (ii) sampling localities. Secondly, we ran partial Mantel tests between venom and genetic (*Dps*) distance matrices. Finally, we tested for correlations between individual-level venom variation and neutral genetic structure using the admixture proportions estimated by TESS as the explanatory variables.

### Venom variation and diet

To test whether geographic variation in venom phenotypes and distribution of toxin genes is associated with differences in diet composition, stomach and gut contents were recorded from 463 preserved, geo-referenced specimens from several museum collections. All prey items were either mammals or reptiles, with the exception of three amphibians, two arthropods and one bird, which were excluded from further analyses to avoid false statistical signals. In total, 327 items were identified at the family level, 249 at the genus, and 192 at the species level.

Frequency of occurrence of prey taxa within localities were used to calculate two metric descriptors of dietary composition at all taxonomic levels: (i) diet niche overlap, describing how similar diet composition is between localities and corresponding to pairwise Bray-Curtis dissimilarity indices; (ii) niche width, calculated using the Shannon diversity index, and describing the diversity of the diet within a locality. Diet composition matrices were used for the Mantel tests. Additionally, we tested for correlations between frequencies of individual prey species and toxin genes in order to identify potential key prey species involved in predator-prey arm races.

### Environmental association analysis (EAA)

One key aim was to test whether the observed variation in venom phenotype and toxin gene distributions were associated with spatial heterogeneity, and hence to identify which environmental factors might contribute to local adaptation and genetic variation.

For climatic and topographic variables, Euclidean distance matrices were calculated based on the average values within each locality, whereas for categorical variables (ecoregion and vegetation) distance matrices were generated based on the proportion of each factor level within localities.

Prior to Mantel test analysis the BIOENV procedure [65] in the *vegan* package was used to reduce the climatic variables to a set of mostly significant. This function calculates Euclidean distances for all possible subsets of scaled climatic variables and finds the maximum Spearman (rank) correlation with the response distance matrix.

In the individual-level analysis, univariate regression models were generated for all variables in order to identify the strength, direction and nature of the relationships between each environmental factor and venom variation/toxin gene presence.

### Gradient analysis

To investigate local environmental patterns at the interface between the two main venom types, we performed a gradient analysis which allows to test associations between phenotypic or genetic variation and environmental factors in a continuous space along a cline. Two roads in Arizona offer the unique opportunity to sample across the transition from venom B to venom A types: one stretching east-west (“Maricopa”) and the other north-south (“Sasabe”). We intensively collected samples from these two transects and tested for presence of MTX and SVMP genes. Trends along the transects were analysed for each climatic variable and correlated with toxin gene presence/absence.

## SUPPLEMENTAL INFORMATION

Supplemental Information includes four figures and four tables.

### ACKNOWLEDGMENTS

We are grateful to the numerous friends and herpetologists who helped with fieldwork and sample collection: D and P. Alvarado, B. & S. Ashley, P.A. Cochran, D. Deem, S.P. Graham, R. Govreau, K. Haas, C.R. Hendry, J.J. Hicks, J. Houck, B. Hughes, T.A. Johnson, R. Legere, C.S. Lieb, P. Lindsey, S.T. Maddock, J.V. Maldonado, C. Meachum, D. & R. Mills, Mrinalini, M. Moffat, E.A. Myers, D. Nixon, D. Ortiz, J.A. Quijada-Mascareñas, J. Rials, D.P. Richards, R. Sawby, G.W. Schuett, B. Starrett, S. Stevens, J. Strickland, T. Tevis, B. Tomberlin, C. Vratil, and D. Weber. G. Whiteley, R. Harrison and R. Morgan performed venom-gland dissection. A. Foote provided insightful comments. Fieldwork carried out under Arizona Game and Fish Department permits SP684786, SP724028 and SP760658 and New Mexico Department of Game and Fish Authorization 3542.

#### Funding

Leverhulme Trust Grant RPG 2013-315 to WW, Santander Early Career Research Scholarship to GZ, Ministerio de Economía y Competitividad Grant BFU2013-42833-P to JJC.

## AUTHOR CONTRIBUTIONS

Conceptualization: WW, GZ; Formal analysis: GZ; Methodology: GZ, JJC, MH; Investigation: all authors; Writing – original draft: GZ; review & editing: all authors.

## DATA AVAILABILITY STATEMENT

The raw reads for the venom-gland RNA-seq experiments have the following accession numbers: *C. scutulatus* type A: XXXXX; *C. scutulatus* type B: XXXX. The raw reads for whole genome sequencing have the following accession numbers: *C. scutulatus* type A: XXXX; *C. scutulatus* type B: XXXX.

